# Photoperiod-dependent developmental reprogramming of the transcriptional response to seawater entry in Atlantic salmon (*Salmo salar*)

**DOI:** 10.1101/2020.03.24.006510

**Authors:** Marianne Iversen, Teshome Mulugeta, Alexander West, Even Jørgensen, Samuel A. M. Martin, Simen Rød Sandve, David Hazlerigg

## Abstract

The developmental transition of juvenile salmon from a freshwater resident morph (parr) to a seawater (SW) migratory morph (smolt) requires a range of physiological adaptations, including the capacity to hypo-osmoregulate. This process, known as smolting, involves both photoperiod-dependent preparative changes before SW is encountered, and activational changes stimulated by exposure to SW. To explore the relationship between these two aspects we undertook experiments in which physiological and transcriptomic responses to SW-challenge were assessed in fish that had experienced different histories of photoperiodic exposure. Compared to fish held on constant light (LL), exposure to short photoperiod (SP) dramatically impaired hypo-osmoregulation in SW, and was associated with extensive glucocorticoid-related changes in gill gene expression. Additionally, a major effect of photoperiodic history was observed in the transcriptional response of LL-acclimated fish to SW, with the response profiles of fish held on LL throughout life being quite distinctive from those of fish which had experienced an 8 week period of exposure to SP prior to return to LL (SPLL). These differences in profile likely reflect a diminishing role for NFAT-mediated responses in SPLL fish, as pathways linked to acute changes in cellular tonicity or intracellular calcium levels decline in importance with preparation for SW.

## 1. Introduction

The gill is the primary site of osmo-sensing and osmoregulatory control in fish (D. H. Evans, Piermarini, & Choe, 2005; T. G. Evans, 2010). In both freshwater (FW) and seawater (SW), osmoregulatory systems work to counter the passive diffusion of ions and water across the gill membranes, and balance plasma osmolality. Euryhaline fish species are defined by their ability to tolerate salinity changes through modulation of osmoregulatory function. In most cases this depends on responses to altered salinity (acclimation), while in a few species groups including salmonids and eels (g. Anguilla), sustained migrations between sea and freshwater are facilitated by preparative changes in osmoregulatory function, forming part of a key developmental life history transition (Folmar & Dickhoff, 1980; Kalujnaia et al., 2007; S. O. Stefansson, Björnsson, Ebbesson, & McCormick, 2008; Jonathan Mark Wilson, Antunes, Bouça, & Coimbra, 2004).

In Atlantic salmon (*Salmo salar*) this preparatory process is commonly known as ‘smoltification’ or, hereafter, ‘smolting’. Smolting is photoperiodically controlled so that migration to sea occurs in a spring ‘smolt window’, when conditions favour juvenile growth (Gross, Coleman, & McDowall, 1988). Smolting requires fish to have previously exceeded a certain size threshold and is presumed to relate to the capacity of juvenile fish to meet the necessary metabolic demands (Higgins, 1985; Kristinsson, Saunders, & Wiggs, 1985; Metcalfe, Huntingford, & Thorpe, 1988; Skilbrei, 1991). During smolting the juvenile salmon develop traits that will enable them to survive in and exploit the marine environment. The increase of photoperiod in spring induces a hormonal cascade influencing behavior, metabolism, growth, pigmentation and gill physiology (Duston & Saunders, 1990; Stephen D. McCormick, 1994; Stephen D. McCormick, Hansen, Quinn, & Saunders, 1998; Stephen D. McCormick, Shrimpton, Moriyama, & Björnsson, 2007). In particular, gill physiology changes in order to accommodate the expected shift in environmental salinity and osmotic drive (D. H. Evans et al., 2005; Kiilerich, Kristiansen, & Madsen, 2007; Nilsen et al., 2007; Pisam, Prunet, Boeuf, & Jrambourg, 1988; Tipsmark et al., 2009). The mitochondria rich cell (MRC), situated on the gill lamella, is a significant component of osmoregulation (Jonathan M. Wilson & Laurent, 2002). The MRC is rich in ion transporters, and change in both morphology and composition in response to salinity (Hiroi & McCormick, 2012; Hwang & Lee, 2007; Hwang, Lee, & Lin, 2011; Madsen, Kiilerich, & Tipsmark, 2009; Pisam et al., 1988). Completion of the smolting process requires entry to sea, where SW exposure triggers the final shifts in physiology and behavior (Lubin, Rourke, & Bradley, 1989; Stephen D. McCormick, Regish, Christensen, & Björnsson, 2013; Nilsen et al., 2007; Pisam et al., 1988). Hence, smolting can be considered a two-step process: a FW preparative phase followed by a SW activational phase.

While the role of photoperiod in timing of preparative changes is well described, less is known about the final changes triggered in smolts during the first few days in SW (Handeland, Berge, Björnsson, Lie, & Stefansson, 2000; Handeland, Jarvi, Ferno, & Stefansson, 1996; Prunet & Boeuf, 1985; S. O. Stefansson et al., 2008), which we will refer to as the SW activational phase. SW responses are also triggered in juveniles entering SW prematurely, which have not initiated or finished the preparative phase of smolt development (Saunders, Henderson, & Harmon, 1985; Stagg, Talbot, Eddy, & Williams, 1989). Triggers may in all cases include osmotic stress due to the hyper-osmotic SW environment as well as direct responses to changes in the concentrations of specific ions (T. G. Evans, 2010; Tyler G. Evans & Somero, 2008; Kültz, 2012). However, the specific response is expected to differ drastically between SW-ready smolts and unprepared juveniles (Houde et al., 2018; Stagg et al., 1989). The importance of SW-exposure for completion of the smolting process and establishment of a SW phenotype is clearly demonstrated by the process of ‘de-smoltification’, which occurs if migration to SW is prevented and involves a loss of tolerance to SW (Arnesen et al., 2003; Sigurd Olav Stefansson, Berge, & Gunnarsson, 1998).

Gill tissue may perceive exposure to SW in at least three possible ways: i) increased cellular tonicity and altered intracellular ion concentrations ii) via cell surface receptors for SW constituents (e.g. Ca^2+^ perceived via the calcium-sensing receptor, CaSR) (Kültz, 2012; Loretz, 2008) and iii) indirectly via hormonal signals (e.g. cortisol, or angiotensin II) which change in response to SW-exposure (Kültz, 2012; Stephen D. McCormick, 2001). In this context, the ‘nuclear factor of activated T-cells’ (NFAT) family of transcription factors have been the focus of recent interest because of their implication in osmo-sensing and in Ca^2+^-dependent transcriptional control (Cheung & Ko, 2013; Hogan, Chen, Nardone, & Rao, 2003; Lorgen, Jorgensen, Jordan, Martin, & Hazlerigg, 2017; Putney, 2012). The NFAT family comprises four subgroups, where groups 1-4 (NFATs c1, c2, c3, c4) are Ca^2+^-stimulated, and the fifth, NFAT5, is regulated in response to extracellular tonicity (Cheung & Ko, 2013; Macian, 2005; Rao, Luo, & Hogan, 1997). All members share a Rel-like homology domain, and bind to similar binding sites in the regulatory region of numerous genes (Macian, 2005).

NFAT5 (also known as osmotic response element binding protein, OREBP, or tonicity-responsive enhancer binding protein, TonEBP), is considered the primordial NFAT, as it is the only one found outside the vertebrate group (Hogan et al., 2003). NFAT5 regulates the transcription of tonicity-responsive genes such as ion transporters and osmo-protective proteins (Cheung & Ko, 2013; Woo, Lee, & Kwon, 2002; Zhou, Ferraris, & Burg, 2006). Hypertonic stress increases nuclear import and retention of NFAT5 through changes in phosphorylation state, while hypotonic stress leads to nuclear export (Cheung & Ko, 2013; Ferraris et al., 2002; Irarrazabal et al., 2010; Macian, 2005).

Two recent studies in salmon focus attention on the role of NFAT signaling during smolting. Lorgen et al. (2015) showed that the salmonid thyroid hormone deiodinase *dio2a* was SW-inducible in gill tissue, and its promoter region was enriched for osmotic response elements (OREs / NFAT5 response elements). A subsequent survey of NFAT5 expression in Atlantic salmon (Lorgen et al., 2017) revealed four NFAT5 paralogues, NFAT5 a1 and a2, and NFAT5 b1 and b2. Of these, NFAT5b1/2 gill expression was highly induced by SW exposure. Together these studies suggest that NFAT5b1/2 could coordinate SW stimulated changes in transcription.

In the present study we sought to extend the previous work on smolting and NFATs to consider the breadth of transcriptional response to SW-exposure in the salmon gill, and to evaluate the extent to which this response relies on NFAT mediated transcriptional control. Our data demonstrate that while NFAT involvement can clearly be seen in the transcriptional response, the importance of this depends to a large degree on the photoperiod to which fish have been acclimated, and the history of prior photoperiodic exposure.

## 2. Materials & Methods

### 2.1 Fish rearing and animal welfare

Atlantic salmon (*Salmo salar*, Linnaeus, 1758) of the Aquagene commercial strain (Trondheim, Norway), hatched and raised (continuous light, LL, >200 lux, 10°C) as part of the ongoing smolt production at Tromsø Aquaculture Research Station (TARS) were used in this experiment. Fish were fed continuously and in excess with pelleted salmon feed (Skretting, Stavanger, Norway).

TARS is approved by the Norwegian Animal Research Authority (NARA) for hold of, and experiments on salmonids, fresh- and salt-water fish and marine invertebrates. When experimental conditions are limited to practices which are undertaken routinely as part of the recognized animal husbandry, with no compromise to welfare, additional formal approval of the experimental protocol by NARA is not required. This is in accordance with Norwegian and European legislation on animal research.

### 2.2 Experimental set-up

The experimental design is presented in fig. 1A.

**Figure 1.**
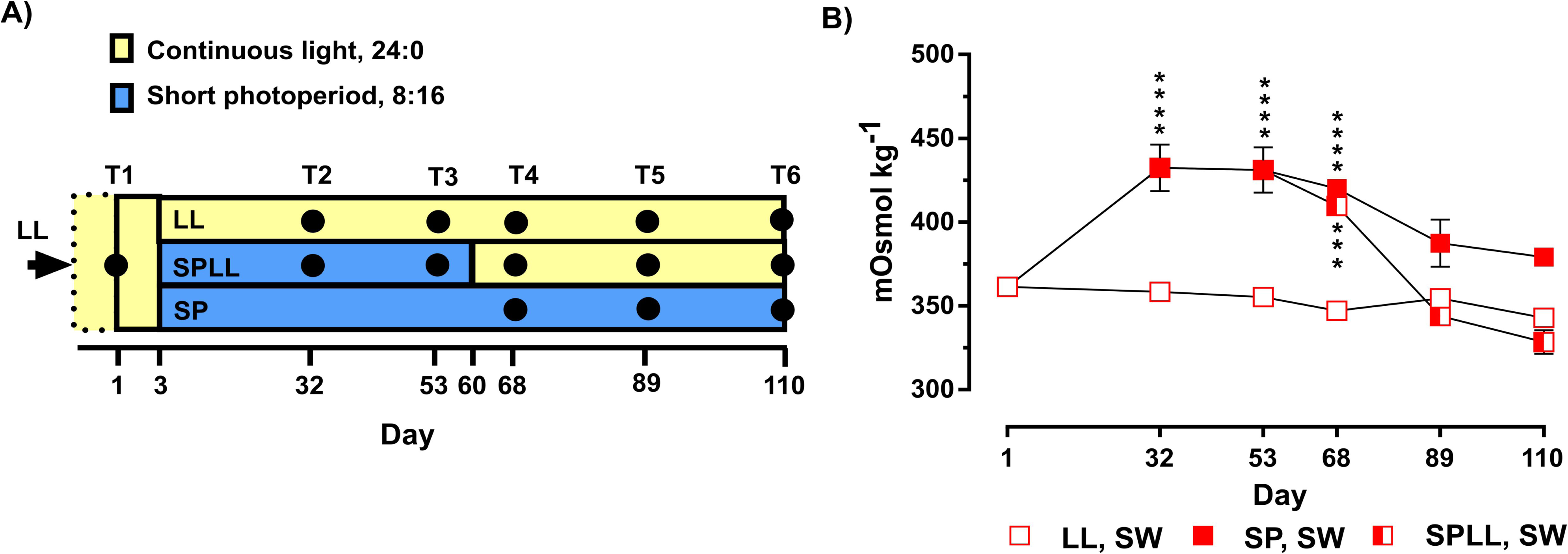
Experimental design and the changing effect of SW challenge on hypo-osmoregulatory capacity. A) Experimental set-up showing the three photoperiod treatments. Samplings are indicated by black dots. B) Plasma osmolality following 24-h SW challenge, data are mean ± S.E.M of n = 6 observations; *** / **** = significantly higher osmolality than at T1, p<0.001 / 0.0001, respectively.

Juvenile salmon, kept in a 500 l circular tank since start of feeding, and at approximately 7 months of age were used for this experiment. A baseline sampling was performed on day 1 of the experiment (mean weight 49.5 g, s.d. ± 7.0 g, n=6); this is referred to as pre-SP. On day 3, 225 juvenile salmon were taken from the original tank and randomly split into two groups of 75 and 150 fish, which were placed in two 100 l circular tanks in separate rooms (FW, 8.5°C). The group of 75 fish were kept on LL for the rest of the experiment. For the group of 150 fish, photoperiod was incrementally reduced from LL to SP (8-h light/16-h dark). Both groups were sampled on day 32 and 53 (n=6 for each treatment). On day 60, half of the remaining fish under SP were moved to a new 100 l circular tank and returned to LL (SPLL). All three groups were then sampled on days 68, 89 and 110 (n=6 for each treatment). During the experiment the fish were fed continuously and in excess over the eight hours corresponding to day in the SP treatment group.

At each sampling point a subsample of fish from each of the treatments were put through a 24-h salt-water challenge (SWC, 100 l tanks, 34 ‰, salinity, 7°C, n=6 for each treatment), starting on the day prior to sampling. The fish were not fed during SWC.

### 2.3 Sampling procedure

Fish were netted out from their respective treatments (including SWC fish) in groups of six. Following anesthesia body mass (±0.5 g) and fork length (±0.1 cm) was measured. Blood was drained from the caudal vein into 2mL lithium-heparinized vacutainers (BD vacutainers®, Puls Norge, Moss, Norway), and placed on ice until further processing. This was followed by decapitation. The operculum was removed from the right side of the head (caudal view), and a gill arch dissected out. The primary gill filaments were cut from the arch and placed in RNAlater® (Sigma-Aldrich, St. Louis, Missouri, USA) for later processing. Samples were stored at 4 °C for 24 h, and then kept frozen at −80°C until further processing.

Blood samples were centrifuged at 6000 x *g* for 10 minutes, and the plasma fraction collected. The plasma was stored at −20°C until later analysis of osmolality could take place. Thawed plasma samples were analysed for osmolyte content using a Fiske One-Ten Osmometer (Fiske Associates, Massachusetts, USA, ± 4 mOsm kg^-1^).

### 2.4 RNA extractions and sequencing

Total RNA was extracted applying a TRIzol-based method following the recommended protocol from the manufacturer (Invitrogen, Thermo Fisher, Waltham, Massachusetts, USA). A NanoDrop spectrophotometer (NanoDrop Technologies, Wilmington, Delaware, USA) was used to check RNA concentration and quality. RNA integrity was confirmed using the Agilent 2100 Bioanalyzer (Agilent Technologies, Santa Clara, CA, USA). RNA was frozen at −80°C until further analysis.

Sequencing libraries (n=167) were prepared with the TruSeq Stranded mRNA HS kit (Illumina, San Diego, California, USA). The 2100 Bioanalyzer using the DNA 1000 kit (Agilent Technologies, Santa Clara, California, USA) was used to determine mean library length, while the Qbit BR kit (Thermo Scientific, Waltham, Massachusetts, USA) was used to determine library concentrations. Samples were barcoded using Illumina unique indexes. Single-end 100 bp sequencing of samples was carried out at the Norwegian Sequencing Centre (University of Oslo, Oslo, Norway), using an Illumina HiSeq 2500.

Removal of sequencing adapters and short sequencing reads (parameters –q 20 –O 8 – minimum-length 40), and trimming of low-quality bases were done using Cutadapt (ver. 1.8.1) (Martin, 2011). Quality control was performed with FastQC software (Andrews, 2010; Andrews, Lindenbaum, Howard, & Ewels, 2011-2014). Mapping of reads onto the reference genome was performed with STAR software (ver. 2.4.2a) (Dobin et al., 2013). Read counts for annotated genes was generated with HTSEQ-count software (ver. 0.6.1p1) (Anders, Pyl, & Huber, 2015). All sequences have been deposited in Array Express, EBI under accession number E-MTAB-8276.

### 2.5 Transcriptome analysis

All transcriptome analysis were performed in R (ver. 3.4.2), using RStudio (ver. 1.0.153).

In order to identify genes that were differentially expressed between the FW and SW sampled fish in the three different treatment groups over the three later time points the R-package EdgeR (ver. 3.14.0) was applied. Raw counts were filtered (expression threshold CPM>1 in five or more libraries), and scaled applying trimmed means of M-values (TMM) scaling. A quasi-likelihood negative binomial generalized log-linear model was used to fit the data, and nine empirical Bayes F-tests were run contrasting between the FW and SW sampled fish for each condition for days 68, 89 and 110 (T4.LL.SW-T4.LL.FW, T4.SP.SW-T4.SP.FW, T4.SPLL.SW-T4.SPLL.FW, etc.). Outputs were filtered requiring a false discovery rate (FDR) of 0.01, and a log_2_-fold change of |1|.

Principal component analysis (PCA) was performed on the full transcriptome using The R Stats Package (stats, ver. 3.4.2) (Love, Huber, & Anders, 2014). Only the three latter sampling points (days 68, 89 and 110) were included in the PCA. For simplicity and interpretability of the plot, TMM normalized counts for each gene in each sample group (n=6, except for T4 SPLL FW where n=5) were averaged before generating the PCA plot.

Lists of differentially expressed genes (DEGs) from each of the sampling groups were compared across time within treatments, and between treatments at the same time point. The numbers of unique and shared DEGs are summarized in the ‘Upset’-plots (UpSetR ver. 1.4.0) (Conway, Lex, & Gehlenborg, 2017) in fig. 2.

**Figure 2.**
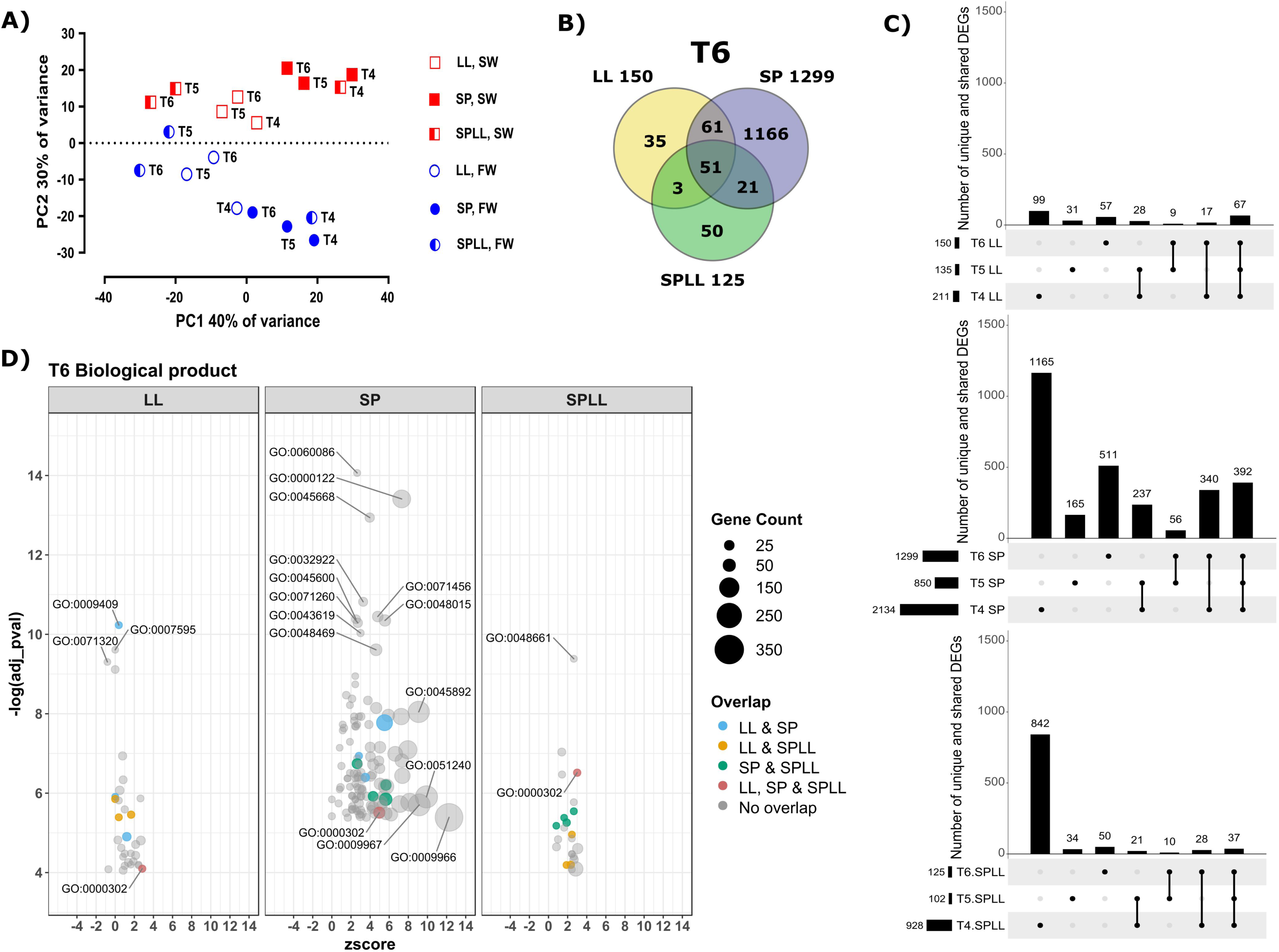
Effect of photoperiodic history on the gill transcriptomic response to SW-challenge. A) PCA plot based on gene expression of the sampled fish. Blue indicates fish sampled from FW and red indicates fish sampled after a 24-h SW challenge. B) Venn diagram showing the number of DEGs (p<0.01, log_2_-FC>|1|) found for each treatment condition at day 110 (T6), and the degree of overlap between the treatments. C) ‘Upset’-plots, indicating how the number of DEGs changed across the three latter timepoints of the experiment for each of the treatments. The bar graph shows number of unique or shared genes for the treatment group(s) indicated by the table below. D) GO-term analysis of SW-sensitive gene expression at T6 for the 3 photoperiod treatments; data are shown as Bubble-plots of enriched biological process (BP) GO-terms and the number of genes linked to each term. Terms enriched across groups are indicated by color. Strongly represented GO-terms are labeled. See supplemental figure S1 for other timepoints and GO categories, and supplemental table S2 for a table of GO-terms and names.

A gene ontology analysis was performed on the same DEG-lists, using topGO (ver. 2.24.0) and the annotation package for the salmon genome Ssa.RefSeq.db (ver. 1.2), with a gill specific gene universe. Fisher statistics and the ‘elim’-algorithm (Alexa, Rahnenführer, & Lengauer, 2006) was applied, with a significance threshold of p<0.05 for enrichment. Only the top 150 GO terms were included in the output. Vizualisation of the GO enrichment using GOplot (ver. 1.0.2) (Walter, Sánchez-Cabo, & Ricote, 2015) and ggplot2 (ver. 3.0.0). GOplot was used to generate the plotting object and z-scores for each GO term (eq.1) that indicate if the trend is towards up- or downregulation of the specific term. The sign of the log2-fold score defines the direction of regulation for each gene. Before plotting unique GO IDs were filtered for a count>5. R-package ggplot2 (ver. 3.0.0) was used for plotting the GO plots, setting a threshold where adjusted p-value <0.0001, or the number of genes annotated to that term >150 for labelling terms in the plot.

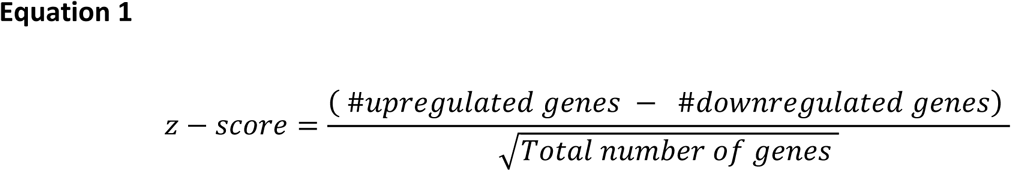

From the set of expressed genes (CPM > 1 in five or more libraries), 18 genes could be identified as NFAT (5), NFAT-like (12) or NFAT-interacting genes (1) based on their SalmoBase annotation (ICSASG_v2) (Lien et al., 2016; Samy et al., 2017). Raw count data was used to calculate mean gene expression at each sampling point for all three treatments. The gene expression of the SP treatment group was then hierarchically clustered using the R-package pheatmap (ver. 1.0.10) (row scaled by z-scores, applying Euclidian distance measures and complete linkage clustering). The resulting order and clustering of genes was then forced onto heatmap of the LL and SPLL groups in order to produce figure 3.

**Figure 3.**
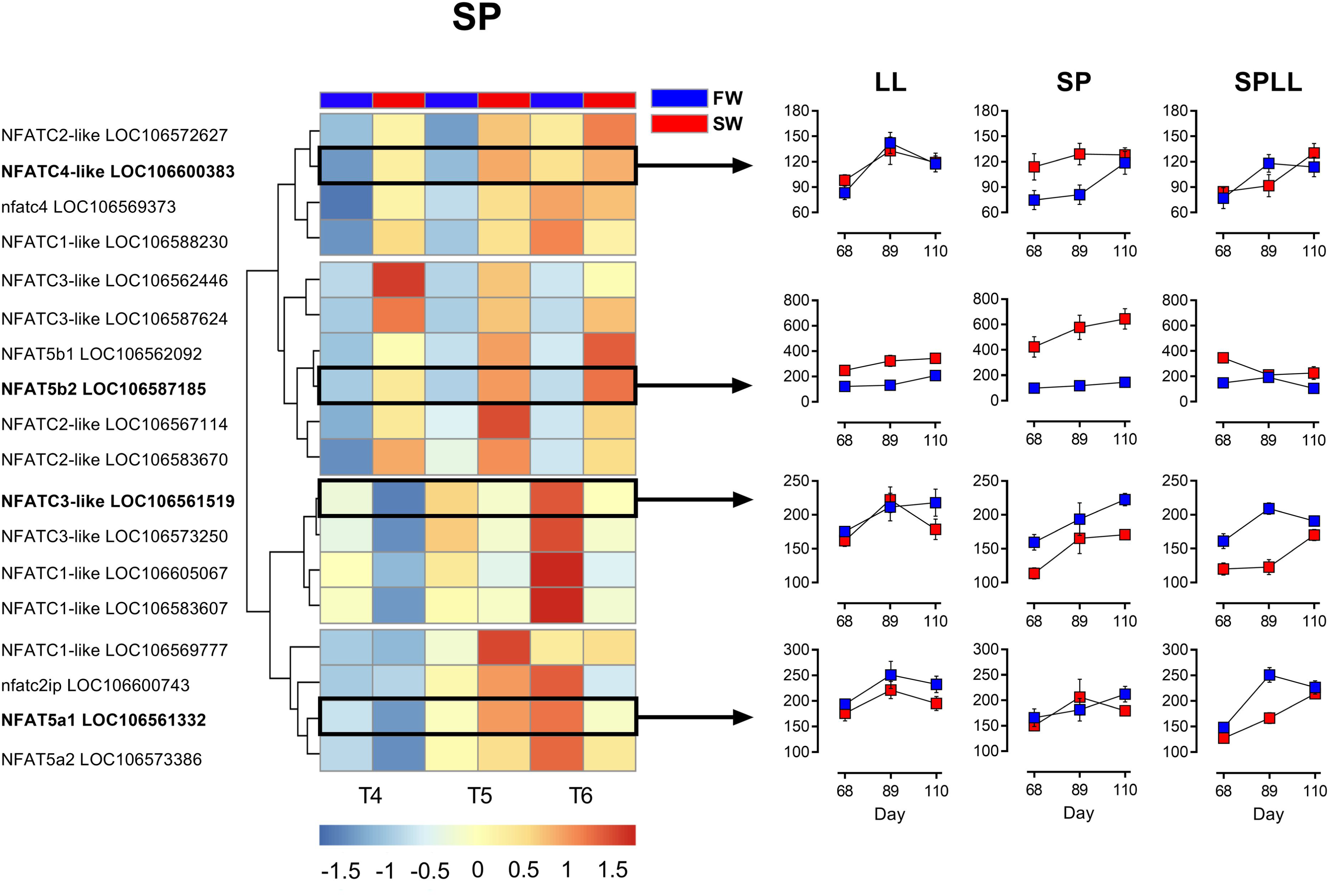
Photoperiodic history-dependent responses of NFAT family members to SW-challenge. The heatmap shows the expression of NFAT-genes (CPM) across the three latter timepoints of the SP-treatment, and graphs on the right show representative profiles of selected NFAT-genes in the 3 photoperiod treatments.

### 2.6 Motif analysis

Motif enrichment analysis was performed using SalMotifDB-shiny tool (https://cigene.no/tools/) (Mulugeta et al., 2019). This tool accesses a database containing over 19,000 predicted transcription factor binding sites (TFBSs) found in the proximal promoter regions (−1,000/+200bp from TSS) of salmonid genes. We used the motif enrichment analysis utility of this tool to screen for enrichment of NFAT and glucocorticoid response element (GRE) motifs in lists of DEGs at the T6 sampling point.

### 2.7 Datasharing

All relevant data can be found within the MS and its supporting information, further the full transcriptomics dataset is accessible in the ArrayExpress depository, with accession number E-MTAB-8276.

## 3. Results

### 3.1 Hypoosmoregulatory capacity

Throughout the study (fig. 1A), fish held on LL upheld the capacity to maintain undisturbed plasma osmolality levels during a 24-h SW challenge (no significant effect of time, P > 0.05, 1-way ANOVA) (fig. 1B). In fish maintained on SP, hypo-osmoregulatory capacity was initially reduced compared to day 1 in LL (P<0.0001 by t-test). As SP exposure extended beyond 8 weeks these fish underwent a partial recovery of hypo-osmoregulatory capacity. Fish that were returned to LL after 8 weeks of SP exposure (SPLL) regained their osmoregulatory capacity within the first four weeks (fig. 1B). Plasma osmolality values of the SPLL group after one week back on LL (T4) were similar to values in SP control fish at the same time point, after which a dramatic improvement in hypo-osmoregulatory capacity was observed (fig. 1B). Eight weeks after return to LL (T6), plasma osmolality values of the SPLL group were 4.2 % lower than in corresponding LL control fish and 9.1 % lower than day 1 values.

### 3.2 RNA profile of the gill response to SW-challenge

To explore treatment effects on the overall RNA expression profile of the gills we performed a PCA analysis (fig. 2A). The first component separated samples by photoperiodic history and sampling time (40% variation explained, PC1) while the second component separated the FW from the SW-challenged fish (30% variation explained, PC2). On the PC1 axis the largest separation of data points was between early (T4, one week after re-entry to LL) and late (T5 and T6, 4 and 8 weeks after re-entering LL) sampling points for SPLL fish. This contrasted with low PC1 resolution for samples from fish in either the LL or SP control groups. The PC2 separation was most pronounced in SP control fish and less so in LL control fish. For the SP and LL groups divergence along PC2 appear independent of time. Contrastingly, in SPLL fish, PC2 resolution was dependent on time of sampling with major segregation between FW and SW samples at T4, one week after re-entering LL, while at both later time points resolution between FW and SW samples was greatly reduced. Overall the PCA analysis indicates that return to LL after SP exposure triggers changes in the gill transcriptome which mirror the improved hypo-osmoregulatory efficiency.

To further investigate the effect of photoperiodic history on SW-responsiveness, we compared lists of SW-DEGs (FDR < 0.01, fold-change > |1|, supplemental material S1) for the 3 photoperiod groups (Fig. 2B, C,). At the end of the study (T6) we found some 10-fold more SW-DEGs in SP fish than in either the LL or SPLL groups. Separate gene ontology enrichment tests were performed for genes responding to SW exposure at T6 in the three photoperiod treatments (supplemental material S3 through S6). Enriched ontologies for SP fish included up-regulated transcripts associated with chromatin silencing and suppression of transcription (e.g. histone deactylase 5, transcriptional repressor p66, NFAT5; GO:0000122 ‘*negative regulation of transcription by RNA polymerase 2*’), and also with formation of stress granules, indicative of translational arrest due to cellular stress (Anderson & Kedersha, 2008) (e.g. ddx6, ddx3x, roquin 1; GO:0010494, ‘*stress granule’*).

Only 51 SW-DEGs (i.e. about 5% of the SP set) were shared across all three photoperiod treatments, and this shared group included genes involved in mitochondrial respiration (e.g. cytochrome P450 subunits, hexokinase-1), presumably reflecting the energy demand imposed by SW challenge. Correspondingly, the only significantly over-represented BP GO-term shared across the photoperiod treatments was GO:0000302, ‘*response to reactive oxygen species’*, encompassing six of the shared genes (fig. 2D).

While there is a similar number of SW-DEGs at T6 in the LL and SPLL treatments (150 and 125 genes, respectively), the overlap between these two groups was almost entirely limited to the universally responsive energy-related genes described above. LL-specific SW-DEGs at T6 were mainly associated with metabolism and cell signaling (f. ex. GO: 0009749 ‘*response to glucose*’, GO:0051591 ‘*response to cAMP*’). In contrast to the SP and LL groups, the SPLL group had a dramatic reduction in DEGs in response to SW between T4 and T6 (Figure 2C). Within the group of SW-induced genes unique to SPLL at the T6 time-point, the inward rectifying K+ channels KCNJ1 and KCNJ5 and ‘junctional cadherin 5 associated’ (JCAD, also know as KIAA1462) were the most strongly induced transcripts (supplemental material S2).

### 3.3 Effects of SW on the expression of NFAT family members

The highly divergent transcriptional responses to SW, including the presence of NFAT5 only in the list of SP-specific DEGs led us to explore further the regulation of expression among all members of the NFAT family of transcription factors (fig. 3, supplemental material S7 and S8). Clustering of response patterns across this gene family gave four distinctive patterns of regulation, represented by the four profile plots in fig 3. The NFAT5b cluster (fig. 3, second cluster from the top) showed strong, SP-specific SW-induction, while weaker SP-specific SW-induction of expression was also seen in the cluster typified by NFAT4c (LOC106600383) (fig. 3, first cluster from the top), but only evident at earlier sampling points (T4, T5). Contrastingly, genes typified by NFAT3c (LOC106561519) showed reduced expression in SW (fig. 3, third cluster from the top). The last cluster of genes were largely SW-unresponsive across the study as a whole (fig. 3, fourth cluster from the top).

### 3.4 Enrichment for NFAT- and GRE-response motifs in SW-DEGs

We used MotifDb ((Mulugeta et al., 2019) (https://salmobase.org/apps/SalMotifDB/) to determine how NFAT response elements are associated with SW-induced changes in gene expression (fig. 4A), focusing on changes occurring at the last sampling point (T6, day 110) of the experiment. This revealed enrichment of seven non-redundant motifs, of which four are associated with SW-induced gene expression changes, in the LL control fish (p<=0.001). Three response elements were enriched in the SP control fish. No enrichment of NFAT elements was seen in SPLL fish at this sampling point. We also looked at presence of glucocorticoid receptor response elements (GREs, fig. 4B) due to the stress response indicated by GO-terms in the SP group, and confirmed that these were only enriched among the SW-response genes in the SP-group (fig. 4B).

**Figure 4.**
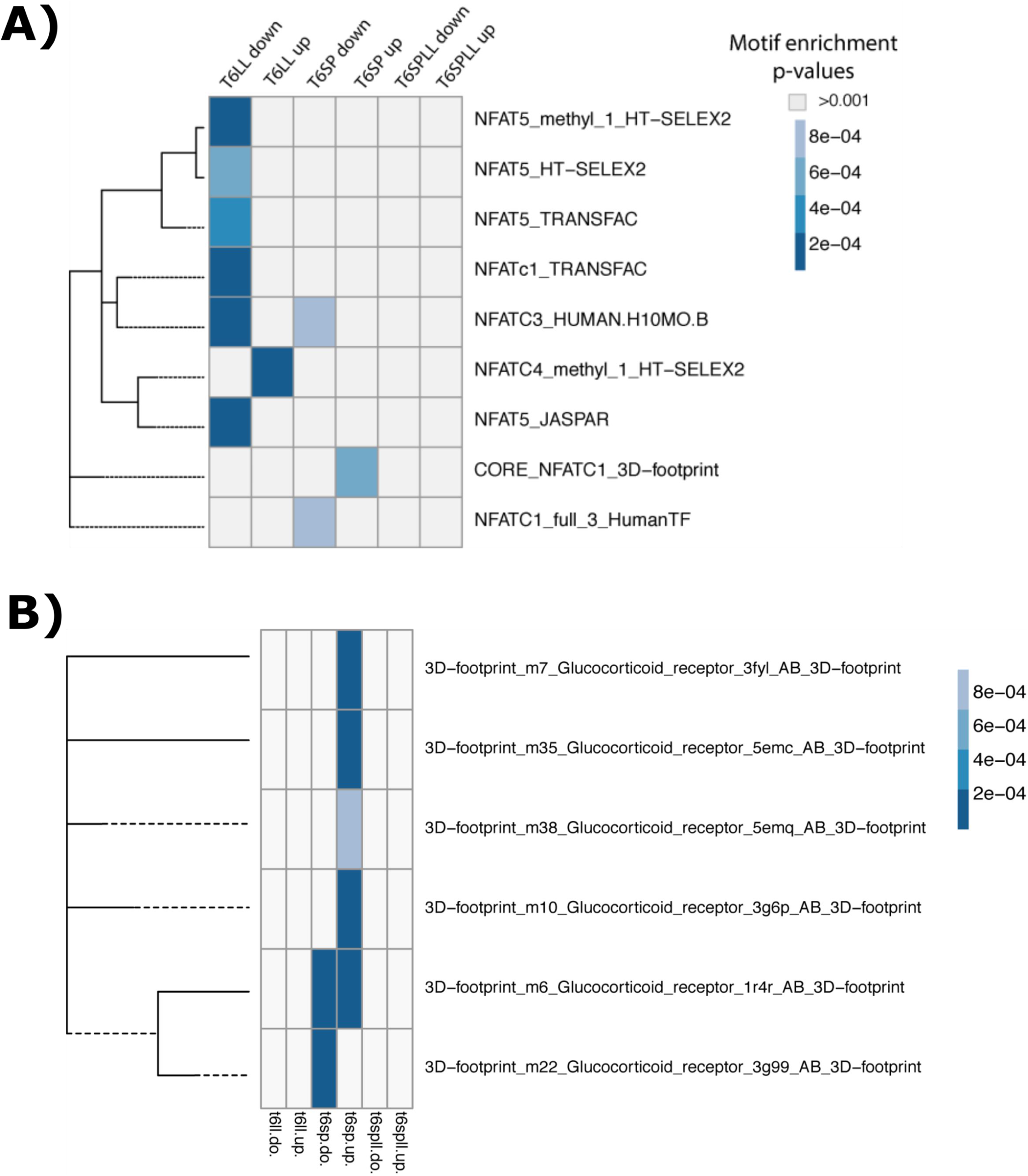
Photoperiodic history-dependent promoter motif enrichment for NFAT and glucocorticoid response elements in SW-induced transcript profiles. Panels A and B show the enrichment of NFAT- and GRE-transcription motifs, respectively, in up- and down-regulated genes at T6, for the three different photoperiod-treatments.

## 4. Discussion

The present study characterizes the effect of photoperiod (SP vs LL) and photoperiodic history (SPLL vs LL) on the gill response to SW exposure in juvenile Atlantic salmon. SP exposure dramatically impairs the ability of juvenile salmon to hypo-osmoregulate in SW and is associated with extensive changes in gill gene expression (fig. 2), including genes predicted to be regulated by the glucocorticoid pathway (fig. 4B), indicative of cellular stress. Contrastingly, exposure of LL acclimated fish to SW does not result in osmoregulatory failure over 24-h, and is associated with less extensive changes in gill gene expression (fig. 2). Nevertheless, a major effect of photoperiodic history was observed in the transcriptional response of LL acclimated fish to SW, with the response profiles of fish held on LL throughout life being quite distinctive from those fish which had experienced an 8 week period of exposure to SP prior to return to LL. The diminished role of NFAT transcriptional regulation in the SW response of SPLL fish observed through reduced motif enrichment analysis (fig. 4A) suggests that preparative effects of SP exposure reduce the involvement of pathways linked to changes in cellular tonicity or intracellular calcium levels in the response to SW.

Previous work by Lorgen et al. (2015; 2017) showed that in the gill the SW-induced gene *dio2a* is enriched for NFAT5 response-elements, and that expression of both dio2a and NFAT5b is SW-induced in SP-acclimated Atlantic salmon juveniles. Our RNAseq analysis confirms these findings, showing that strongest SW-induction of NFAT5b is indeed seen in SP acclimated fish, as well as implicating NFAT4 and NFATc3 in the response. Given that this is the case, it is somewhat surprising that statistical enrichment for NFAT motifs is less pronounced within the SW-induced transcriptome of SP fish than in LL fish. We believe this may reflect a swamping of signal by large numbers of genes induced through stress-activated pathways, including but probably not limited to the adrenal corticoid axis revealed by GRE enrichment in SW-induced genes in SP fish. In support of this interpretation the subset of SW-induced genes shared between fish in the LL and SP T6 groups, which constitutes less than 10% of the overall SP SW-induced group (but about half of the LL SW-induced group) is highly enriched for NFAT5 elements (p<0.01).

Despite the superficial similarity observed between the LL and SPLL fish in ability to hypo-osmoregulate (fig. 1B) as well as the magnitude of transcriptional responses to SW exposure (fig. 2), it is clear from the GO analysis that the SW-responses of fish in these two groups are quite distinctive. We suggest that the marked enrichment of NFAT-response elements, and in particular NFAT5, in the LL group reflects a transient activation of NFAT5-responsive genes in response to SW. By contrast, in the SPLL group there is no motif enrichment for NFAT5 nor the Ca^2+^-regulated NFATs. We interpret this lack of NFAT5 responses in SPLL as evidence for NFAT5-signaling playing a role in the activation of hypo-osmoregulation in salmon which have not developed a SW migratory phenotype. Accordingly, exposure to SP for 8 weeks prior to re-exposure to LL stimulates pre-adaptation and obviates the need for NFAT-mediated responses to SW exposure – presumably because even in the initial phase of SW exposure, pre-adapted gill cells do not experience significant changes in tonicity or intracellular Ca^2+^ levels.

The transcriptional response of the NFAT family was not limited to NFAT5b since we also observed SW-induction of NFATc1 and c4 in the SP group, and photoperiodic history-dependent SW-suppression of NFATc3 and NFATc1 paralogous pairs in the SP and SPLL groups. In mammals, these calcium-regulated NFAT’s play important roles in immune function, but also in the development, differentiation and function of various other cell types such as osteoclast and cardiac tissue (Ames, Valdor, Abe, & Macian, 2016; Hogan et al., 2003; Macian, 2005). Changes in intracellular calcium leading to NFAT activation may conceivably arise as a result of Ca^2+^ production as a second messenger within the cell, or as a result of Ca^2+^ entry from the environment – and both these pathways are likely to be involved in osmosensing (Kültz, 2012).

In addition, extracellular Ca^2+^ may affect gill function through the G-protein coupled calcium sensing receptor (CaSR), expressed in the MRCs and proposed to function as a salinity sensor in fish (Loretz, 2008; Loretz, Pollina, Hyodo, & Takei, 2009; Nearing et al., 2002). While CaSR signal transduction has primarily been linked to cAMP-dependent signal transduction, the possibility of cross-talk with NFAT pathways is suggested by work on TNF secretion in the mammalian kidney tubule (Abdullah et al., 2006; Gong & Hou, 2014).

Our results clearly show that NFATs are playing a minor role in SW regulated transcriptional responses in SPLL fish compared to LL and SP. This is consistent with a model where the photoperiodic treatment received (SPLL) is known to stimulate a range of smolt characteristics including improved long-term performance in SW (Berge et al., 1995; S. D. McCormick et al., 1995; Stephen D. McCormick et al., 2007; Saunders et al., 1985; S. O. Stefansson et al., 1991; S. O. Stefansson et al., 2008). With the exception of day 68 (i.e. the first week after return to LL from SP, when these fish are in a transitional state), there is no SW-induction of NFAT5b-expression or any other NFATs, nor is there any enrichment of NFAT-motifs in the SW-responsive transcriptome. Nevertheless, a small number of genes were uniquely stimulated by SW in the SPLL group. These included the inward rectifying potassium channel genes KCNJ1 and KCNJ5, the former being ATP-regulated and the latter being G-protein regulated (Clapham, 1994; Ho et al., 1993; Krapivinsky et al., 1995). Also, we find the cardiac regulatory gene junctional protein associated with coronary artery disease, known as JCAD. The potassium channels have been identified as key markers for SW adaptation in eels, where they have been found to be expressed in MRCs (Suzuki et al., 1999; Tse, Au, & Wong, 2006). JCAD is predicted to play a role in endothelial cell junctions (Akashi, Higashi, Masuda, Komori, & Furuse, 2011) and has been linked to the Hippo signaling pathway (Jones et al., 2018), which regulates cell proliferation and apoptosis (Halder & Johnson, 2011). Both KCNJ1 and JCAD show high SW-inducibility after being exposed to the photoperiod-induced smolting (S2), and they therefore represent the final activational response to SW occurring specifically in fish that have completed a FW preparative phase in response to photoperiod. Further studies to understand the impact of these genes on gill function in SW are now warranted.

## Supplemental material

**S1** Overview of DEGs for each condition and timepoint, after filtering for FDR<0.01 and a log_2_-fold change of |1|.

**S2** Boxplots of the genes KCNJ1, KCNJ5 and JCAD, raw counts.

**S3** Additional GO plots showing how GO-enrichment varies over time and between treatments.

**S4, S5 and S6** Results from the GO analysis and overview of the GO-terms that are included in the plots after filtering for number of connected genes and log of the adjusted p-value for enrichment.

**S7** Overview of NFAT-genes, including raw count data. Genes are ordered as in the heatmap.

**S8** Boxplots of the NFAT genes, raw counts.

